# Telomere-to-telomere reference genome of the common five-lined skink, *Plestiodon fasciatus* (Squamata: Scincidae)

**DOI:** 10.1101/2025.07.03.663019

**Authors:** Jon J. Hoffman, Frank T. Burbrink, R. Alexander Pyron, Christopher J. Raxworthy

## Abstract

Although the publication of high-quality reference genomes is steadily increasing, many clades remain chronically neglected. Skinks (order: Squamata; family: Scincidae) are one of the most diverse lizard families (1,785 species), yet there are currently just six published chromosome-level skink genomes. Here, we present the first telomere-to-telomere, chromosome-level reference genome for one of the most abundant lizards in the eastern United States, the common five-lined skink (*Plestiodon fasciatus*). Through the sequencing of RNA, long-read DNA, and Hi-C chromatin interactions, we produced an annotated reference genome (N50 = 227MB, L50 = 3) consisting of 6 macrochromosome pairs and 7 microchromosome pairs with 98% of BUSCO genes represented (lineage: Sauropsida; 7480 BUSCO markers) represented, providing one of the most complete skink genomes to date: *rPleFas1.1*. Functional annotation predicts 32,520 protein-coding genes (16,100 unique, named genes) with an average gene length of 9,372bp. Repeat annotations estimate that transposable elements comprise 46.7% of the genome, for which we show the amount and content is remarkably conserved across Scincidae.

## Introduction

With the advent of whole-genome sequencing, numerous genomic resources are now available for many clades across the vertebrate tree of life, with birds and mammals comprising most of these genomes. Despite the abundance of ecological and evolutionary research on squamate reptiles, the availability of high-quality genomes has, until recently, lagged far behind other amniote groups. There has been a recent drastic push to increase the production of long-read genomes for many squamate groups (Gable et al., 2023). As a result, the number of reference-level genomes of NCBI has risen from 115 in 2023 to 292 in Spring of 2025, a ∼150% increase over the last year and a half. Still, there is an underrepresentation of diverse families such as skinks, geckos, chameleons, and amphisbaenids (Pinto et al., 2023). Generating high-quality reference genomes across squamates is paramount, as broad sampling is an important aspect of understanding and preserving a genetic record of biodiversity, hypothesizing species relationships, and conservation of endangered taxa (Brandies et al., 2019; Worley et al., 2017)

For example, there is a large variation of chromosomal architecture within squamates, with karyotypes ranging from 2n=16 in the gecko *Gonatodes taniae* to 2n = 62 in the microteiid *Notobrachia ablephara* and the chameleon *Rieppeleon brevicaudatus* (Mezzasalma et al., 2024; Pellegrino et al., 1999; Rovatsos et al., 2017; Schmid et al., 1994). In skinks, the number of chromosomes ranges from 2n=22 to 2n=32, and genomic architecture is relatively conserved (Deweese and Wright, 1970; Giovannotti et al., 2009). Most *Plestiodon* have 13 chromosome pairs, where 2n=26 with 6 macrochromosome pairs and 7 microchromosome pairs (Xu and Zhu, 2024), though *P. anthracinus* has 12 chromosome pairs (Hardy et al., 2017).

There is also great diversity of sex-determination systems in squamates (Alam et al., 2018; Ezaz et al., 2009; Janzen and Phillips, 2006), ranging from temperature-dependent sex determination to genotypic sex determination with XY/XX (male heterogamy) and ZZ/ZW (female heterogamy). Sex-determining systems are extremely labile in squamates (Ezaz et al., 2009; Mezzasalma et al., 2021). For example, it is estimated that there have been 17–25 in sex-determination transitions in geckos, with multiple sex-determining systems in some genera (Gamble et al., 2015). Unlike geckos but similar to iguanas, skinks have a conserved XY sex determining system (male heterogamy) with homomorphic sex chromosomes that are difficult to distinguish (Kostmann et al., 2021; Xu et al., 2024). There is little apparent variation in the sex-determining system of skinks, but this may be an artifact of limited genomic resources (Dodge et al., 2023). It has been hypothesized that the XY sex-determination system may have evolved independently from that of other squamates, such as *Podarcis mualris* (Lacertidae) and *Anolis carolinensis* (Dactyloidae), which are commonly used in comparative studies (Kostmann et al., 2021).

Skinks represent 4% of amniote diversity, yet there are currently only six chromosome-level skink genomes, four of which are Australian. To increase both the phylogenetic and biogeographic diversity of genomic resources for skinks, we present a telomere-to-telomere annotated reference genome for *Plestiodon fasciatus*. The common five-lined skink, *P. fasciatus* (Linnaeus 1758), is one of the most abundant lizards of the eastern U.S and southeastern Canada (Powell et al., 2016). These generalist skinks are small (total length: 12.5–22.2 cm, maximum snout–vent length: 8.6 cm), found in mesic wooded areas, and reside in cover materials, such as rock crags, logs, and tree bark, but will emerge to thermoregulate or search for invertebrate prey (Brazeau et al., 2015; Fitch and Fitch, 1954; Powell et al., 2016). This genome is the seventh chromosome-level in the family Scincidae, which currently includes *Bassiana duperreyi* (Hanrahan et al., 2025), *Spondylurus nitidus* (Rivera et al., 2024), *Carinascincus ocellatus, Tiliqua scincoides, Cryptoblepharus egeriae* (Dodge et al., 2025), and the congener *P. gilberti* (Richmond et al., 2025). Barring *P. gilberti*, all of the other species represented which shares a common ancestor with *P. fasciatus* ∼115Ma (Title et al., 2024), while *P. gilberti* and *P. fasciatus* diverged ∼17Ma (Brandley et al., 2011)

## Materials & Methods

### Sample Acquisition

We collected an individual female *Plestiodon fasciatus* in Allegan County, Michigan (Lat: 42.5394; Long: -85.9949), and euthanized with MS-222 prior to tissue collection following Michigan DNR permits and approved IACUC protocols at the American Museum of Natural History (AMNH). We sampled liver, lung, heart, skeletal muscle, kidney, and skin from the individual and stored them in NAP buffer (Camacho-Sanchez et al., 2013) to preserve the RNA. Due to a suboptimal PacBio Revio whole-genome sequencing effort of the NAP preserved liver tissue from this individual, we collected a female *P. fasciatus* in McCracken County, Kentucky (Lat: 37.1501; Long: -88.7953), with Kentucky Department of Fish and Wildlife permits to provide a blood sample (stored in EDTA) for genomic sequencing. Despite the 650 km between these localities, there are no major biogeographical barriers (Soltis et al., 2006). Furthermore, there is little mitochondrial phylogeographic structure within the population (Howes and Lougheed, 2008) and therefore likely little genomic variation, though this has not been tested. Both individuals are vouchered specimens, catalogued in the Herpetology Collections at the American Museum of Natural History (AMNH) as AMNH-179334 (Allegan County, MI) and AMNH-179327 (McCracken County, KY).

### RNA extraction and sequencing

For RNA sequencing, we sent six tissues (liver, lung, heart, skeletal muscle, kidney, and skin) to Azenta/Genewiz for extraction, library preparation with poly(A) selection to target eukaryotic strand-specific mRNA, and sequencing on an Illumina NovaSeq 2×150bp, generating ∼100M paired-end reads. All RNA sequences passed the initial quality check run with FastQC (Brown et al., 2017). Sequencing adapters in the resulting sequences were filtered and trimmed with trimmomatic using the default settings (Bolger et al., 2014).

### Genomic DNA extraction and sequencing

Genomic DNA was extracted from blood stored in EDTA from AMNH-179327 using the Qiagen MagAttract High Molecular Weight DNA kit following their ‘Manual Purification of High-Molecular Weight Genomic DNA from Whole Blood’ protocol from the MagAttract HMW DNA Handbook in the AMNH ICG. The extracted DNA was then sent to Azenta/Genewiz, where it was sequenced with PacBio Revio HiFi sequencing on 1 SMRT cell, which typically results in ∼15 million reads and ∼100GB of data, depending on the quality of the input sample. We expected a coverage of ∼66x based on an estimated 1.5GB genome size. Sequencing generated ∼6.3 million read pairs and ∼79GB of data, with an average read length of ∼12,000bp, resulting in ∼53x coverage.

Finally, we generated Hi-C data with Phase Genomics using the Proximo® Hi-C animal genome scaffolding platform from a collected blood sample. Proximity-ligated fragments were sequenced on an Illumina NovaSeq to produce 2×150bp paired-end reads. The sequencing generated 300 million read-pairs, of which 56% were high quality, yielding 1,721,863 read-pairs per contig that were ‘usable’, indicating that they mapped to different >5kb contigs.

### Draft Genome assembly

A draft genome assembly was made from the HiFi long-read sequencing using Hifiasm v0.25.0, a haplotype-resolved de-novo assembly tool for PacBio HiFi reads (Cheng et al., 2021). Hifiasm was run without the Hi-C sequences as its inclusion led to a drastic increase of contigs, from 34 without Hi-C to ∼6,000 with Hi-C. Due to the high content of low-divergence repeats, we soft-masked the draft genome with the *Earlgrey* v4.1.1pipeline (Baril et al., 2024) prior to Hi-C mapping, which also provide transposable element (TEs) annotations (see below).

### Hi-C incorporation

To incorporate the Hi-C sequencing with the draft genome assembly, we initially aligned the Hi-C sequences to the HiFi draft genome with a Burrow-Wheeler Alignment (*BWA* v 0.7.19) (Li, 2013). Then, we processed the resulting alignments with *SAMtools* v1.22.1 (Danecek et al., 2021) to remove duplicate sequences. We then scaffolded the assembly using the *de novo YAHS* v1.2.2 assembly pipeline (Zhou et al., 2023). From there, we used Juicer Tools to generate a Hi-C contact map (Durand et al., 2016). Finally, we visualized the scaffolding of the chromosome-level assembly with *Juicebox Assembly Tools* v2.20.0 (Robinson et al., 2018).

### TE and gene annotation

We annotated the final assembly by first modelling and quantifying TEs by soft-masking the genome with *EarlGrey* v4.1.1, a fully automated TE annotation pipeline (Baril et al., 2024) with the default settings, which includes a 100bp minimum length and 10 iterations of the BLAST, extract, and extend process. Next, we functionally annotated the predicted gene regions using the train, predict, update, fix, and annotate steps of the *funannotate* pipeline (Palmer and Stajich, 2020). The ‘train’ step aligns RNA-seq data, assembles it with *Trinity* (Grabherr et al., 2011), and runs *PASA*, which models gene structures based on alignments of expressed transcripts (Haas et al., 2003). The ‘predict’ step uses *PASA* gene models to train *Augustus*, a *de novo* gene finder (Stanke et al., 2008), prior to running *EvidenceModeler* (Haas et al., 2008). We included RNA-seq data from the liver, lung, heart, skeletal muscle, kidney, and skin as evidence for *EvidenceModeler*. We used the Tetrapoda BUSCO database with *Taeniopygia guttata* as the seed species for the ‘predict’ step and kept the default options for each step. The ‘update’ step fixes gene models that disagree with RNA-seq data, which are corrected in the ‘fix’ step. Prior to running the ‘annotate’ step, we ran *InterProScan* v5.74-105.0 (Jones et al., 2014) to run predicted genes against the InterPro database for gene families and downloaded *Eggnog-mapper* v2.1.12 (Cantalapiedra et al., 2021) locally to be run during the ‘annotate’ step. The ‘annotate’ step incorporates the generated data into an annotated genome. We used the default functional annotation databases in the annotation step. To compare across existing chromosome-level skink genomes, we ran the above pipeline on the assemblies of *Tiliqua scinciodes, Spondylurus nitidus, Bassiana duperreyi, and Carinascincus occelatus*.

### Synteny analysis

To assess genomic synteny of the *Plestiodon fasciatus* genome and other chromosome-level squamate assemblies, we created a custom pipeline called *Synk* (https://github.com/jomhoff/Synk) that uses the output files from *compleasm* (Huang and Li, 2023) and isolates the BUSCO genes to create comparative text files, and uses *RIdeogram* (Hao et al., 2020) to plot the syntenic chromosomal regions from BUSCO genes between species in one script. Compared to methods for calculating and visualizing whole genome synteny, *Synk* runs much faster. In addition, limiting the dataset to BUSCO genes minimizes paralogy issues, as BUSCO genes are highly conserved, single copy orthologs. This is especially effective in ensuring appropriate estimates of synteny in cross-species and cross-genera comparisons. Here, we show synteny between *P. fasciatus, Bassiana duperreyi, Spondylurus nitidus, Carinascincus ocellatus, Tiliqua scincoides*, and *P. gilberti*.

## Results and Discussion

### Assembly of the P. fasciatus genome

After completing *Hifiasm* with the long-read PacBio sequences, the draft assembly of the genome was close to complete, with an L50 of 3 and a total of 34 contigs, 9 of which represented near telomere-to-telomere chromosomes (**Table 1**). With the Hi-C data incorporated, we successfully generated a chromosome-level assembly of *Plestiodon fasciatus* with 18 unplaced scaffolds that we refer to as *rPleFas1*.*1* after Vertebrate Genome Project naming rules (Rhie et al. 2021) (**Table 1; Figure 1**). One of the 18 unplaced scaffolds includes the complete mitochondrial genome. The other 17 unplaced scaffolds range in size from 8,262bp to 1,241,752bp and consist of unplaced TEs and mRNA sequences. For the complete autosomal genome, the presence of telomeres was estimated with the characteristic rise in GC content at the ends of chromosomes (**Figure 1**) and confirmed with *tidk* v0.2.65, a toolkit for identifying telomeres that outputs a count of telomeric repeats in windows across the genome (Brown et al., 2025). There is also a large spike in GC content in the middle of the second chromosome, which corresponds to a repeat-dense region consisting of Long Interspersed Nuclear Elements (LINE), simple, and unclassified repeats.

**Table 1:** Assembly statistics of the genome before and after Hi-C incorporation.

|  | Metric | Draft Genome | Hi-C Informed Genome |
| --- | --- | --- | --- |
| **Basic Stats** | Total assembly size (bp) | 1,537,706,373 | 1,537,706,373 |
|  | Number of contigs | 34 | 34 |
|  | Number of scaffolds | 34 | 31 |
|  | GC content (%) | 45.6 | 45.6 |
| **Contiguity** | Contigs N50 (bp) | 166,278,287 | 166,278,287 |
|  | Contigs L50 (bp) | 3 | 3 |
|  | Largest Contig (bp) | 244,253,184 | 244,253,184 |
|  | Scaffolds N50 (bp) | 166,278,287 | 227,050,938 |
|  | Scaffolds L50 | 3 | 2 |
|  | Largest Scaffold (bp) | 244,253,184 | 304,269,845 |
| **Completeness** | BUSCO Complete (%) | 98.17 | 98.17 |
|  | Missing BUSCOs (%) | 1.24 | 1.24 |
|  | Fragmented BUSCOs (%) | 0.19 | 0.19 |
|  | Sequencing coverage (X) | 53 | 116 |
|  | Telomere to Telomere Chromosomes | 6 | 13 |

**Figure 1:**
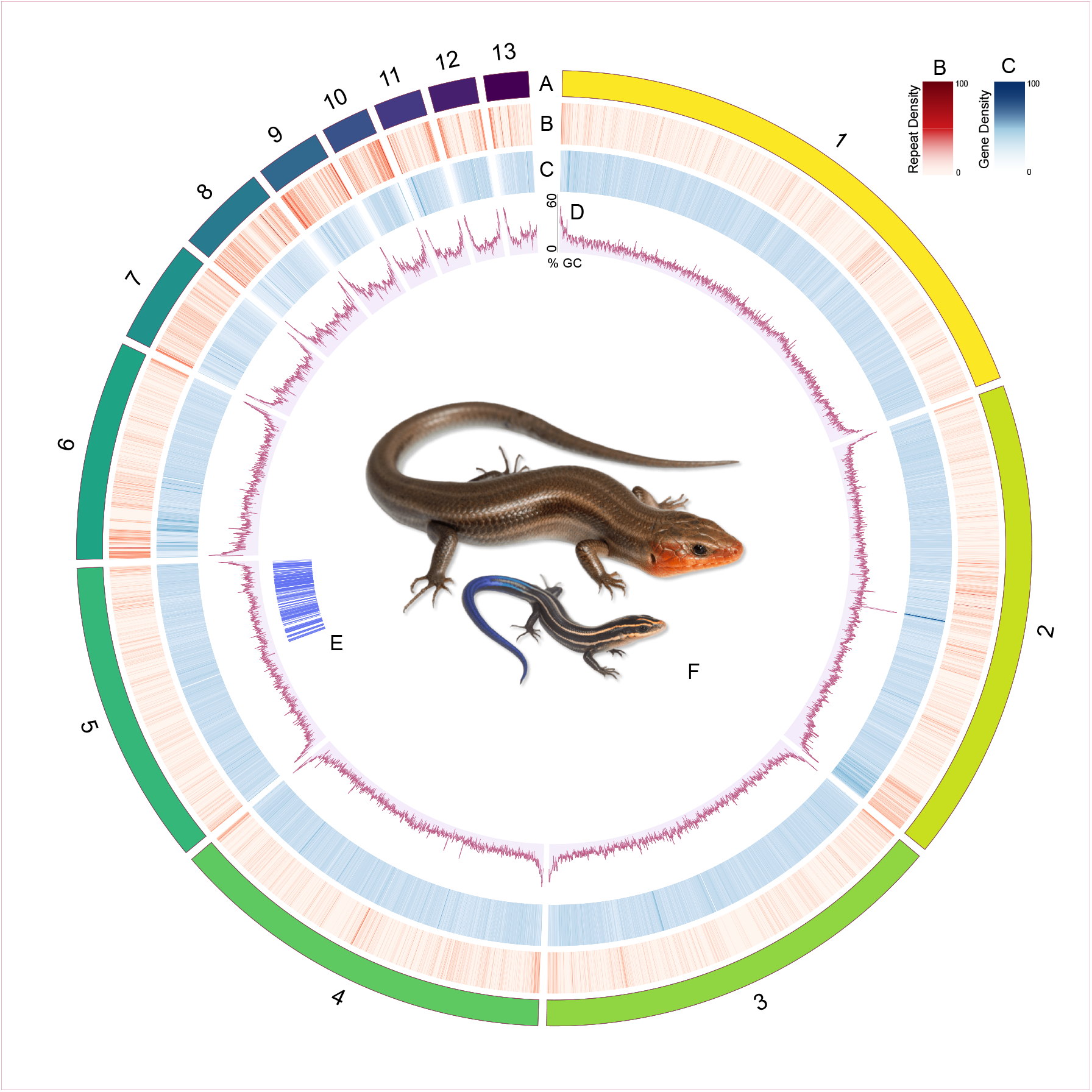
Ideogram for *Plestiodon fasciatus*. **Row A** represents chromosome number and relative size. **Row B** is a heatmap of gene density for 50kb windows, where darker colors represent a higher density of genes. **Row C** is a heatmap of repeat elements for 50kb windows, where darker colors represent a higher density of repeat elements. **Row D** is a line chart of GC content across the genome. **E** is the the region of the genome that is syntenic with the X chromosome of *Bassiani duperreyi*. **F** is a photograph of an adult male and juvenile *Plestiodon fasciatus* (photo via Herps of Arkansas). Plot made with Circos (Krzywinski et al., 2009).

### Annotation of genes and TEs

The resulting annotations from the *funannotate* pipeline consists of an estimated 32,520 genes. Filtering for named genes and removing isoforms resulted in 16,100 unique genes with common names (**Table 2**). Like other squamates, the genome of *P. fasciatus* has a high proportion of repetitive regions which account for 46.7% of the genome (Pasquesi et al., 2018). Unclassified repeats account for 23.3% of the genome, while DNA repeats and LINEs respectively make up 7.0% and 9.3% of the genome (**Figure 2**). The Kimura 2-parameter distance between the repeat sequences produces an approximately bimodal distribution, with a peak indicating a large degree of moderately diverged repeats and a peak indicating recently diverged repeats (**Figure 2**). The more diverged peak is mainly driven by unclassified repeats. Interestingly, this a pattern that is seen in all five skinks tested. Furthermore, the more recently diverged peak is driven by a large proportion of both DNA and LINE repeats and is also conserved across the species analyzed (**Figure 2**).

**Table 2:** Annotation Statistics of the Genome after the *Funannotate* pipeline.

| Category | Metric | Value |
| --- | --- | --- |
| Gene Content | Number of protein-coding genes | 32,520 |
|  | Unique genes with common names | 16,100 |
| Gene Structure | Average gene length (bp) | 9,372.34 |
|  | Total number of exons | 252,300 |
|  | Average exon length (bp) | 255.27 |
|  | Multiple-exon transcripts | 25,572 |
|  | Single-exon transcripts | 7,372 |
| Functional Annotation | Genes with GO term annotations | 21,578 |
|  | Genes with InterProScan hits | 24,316 |
|  | Genes with eggNOG annotations | 26,007 |
|  | Genes with Pfam domains | 19,449 |
|  | CAZyme-annotated genes | 319 |
|  | MEROPS-annotated genes | 1,143 |

**Figure 2:**
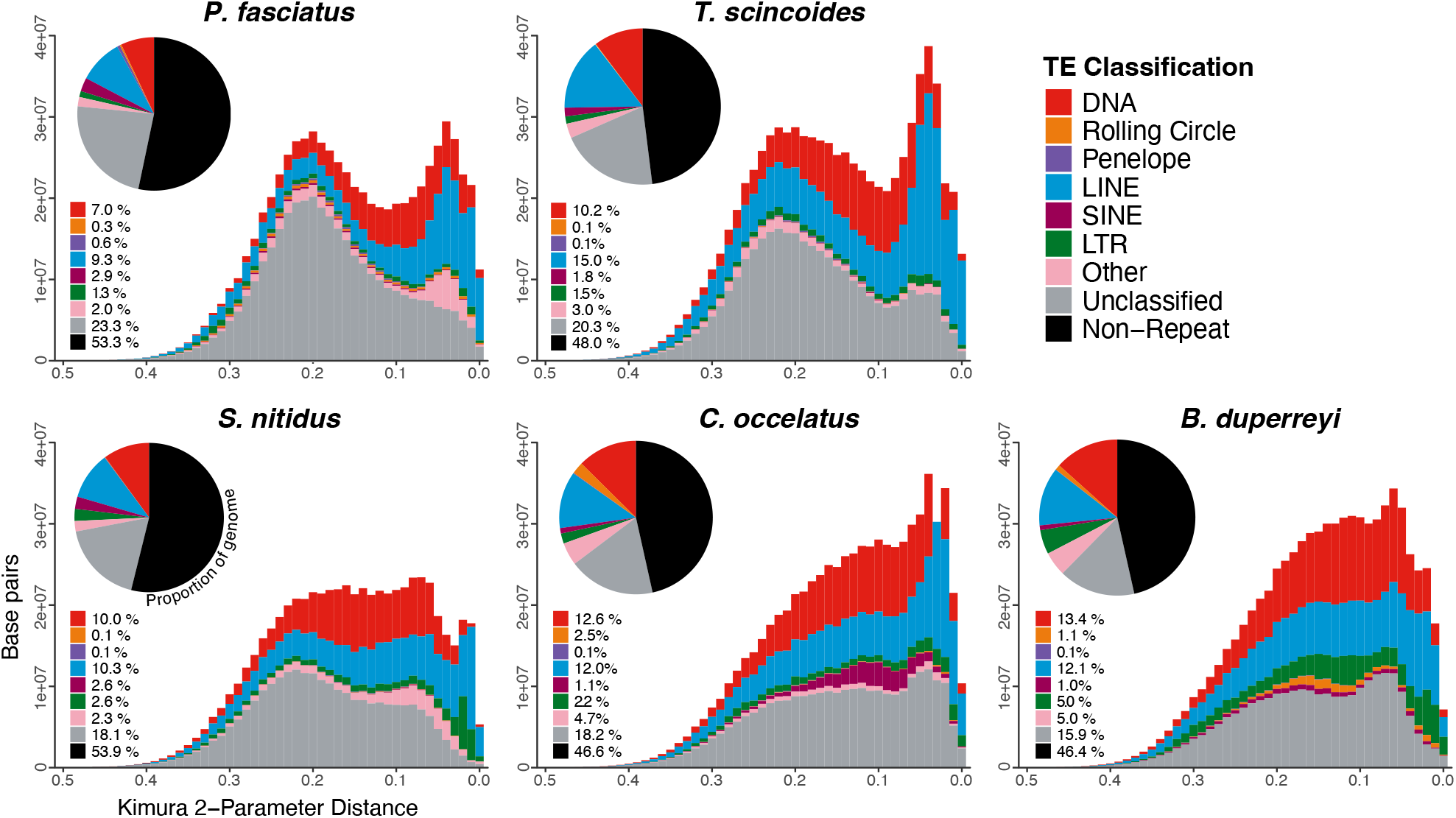
Divergence estimates of TEs shown with Kimura 2-Parameter Distance for five chromosome-level skink genomes where the higher the Kimura 2-Parameter Distance, the more diverged the sequences. The proportion of each transposable element subdivision in the genome for each species is displayed with a pie chart.

Although the impact of TEs on the diversification and adaptation of squamates has yet to be studied in detail, it has been hypothesized that variation in TEs can impact phenotypic adaptation through TE domestication, exaptation, host-gene regulation, formation of retrogenes, and genomic plasticity (reviewed in Schrader and Schmitz, 2019), and that TEs contribute significantly to genomic variation (Catlin and Josephs, 2022). Since TEs are highly mobile across the genome, are abundant, labile, and have been major players in the evolution of eukaryotic genomes (Bowen and Jordan, 2002; Oliver and Greene, 2009), we expect variation of TEs among distantly diverged clades. Despite originating around ∼115mya, the skinks in this study display remarkable conservation of TE composition, especially when considering the variation seen in younger clades, such as extant mammals (Platt et al., 2018) and plethodontid salamanders (Sun et al., 2012).

### Synteny of skink genomes

The genomes of *Plestiodon gilberti* and *P. fasciatus* appear highly syntenic, with the only rearrangement being an inversion in the eighth chromosome. Despite the ∼150ma divergence (Title et al., 2024), the macrochromosomes are largely conserved across the skink species analyzed, aside from a large inversion on chromosome 1 between *Tiliqua scincoides* and all other species (**Figure 3**). Compared to the other scincids, *P. fasciatus* and *P. gilberti* have fewer microchromosomes, where chromosomes 14, 15, and 16 in *S. nitidus* and *T. scincoides* are syntenic with chromosome 8 in *P. fasciatus* and *P. gilberti*. There is also a rearrangement creating syntenic blocks relating chromosomes 6 and 7 in *S. nitidus* and *T. scincoides* to chromosome 5 in

**Figure 3:**
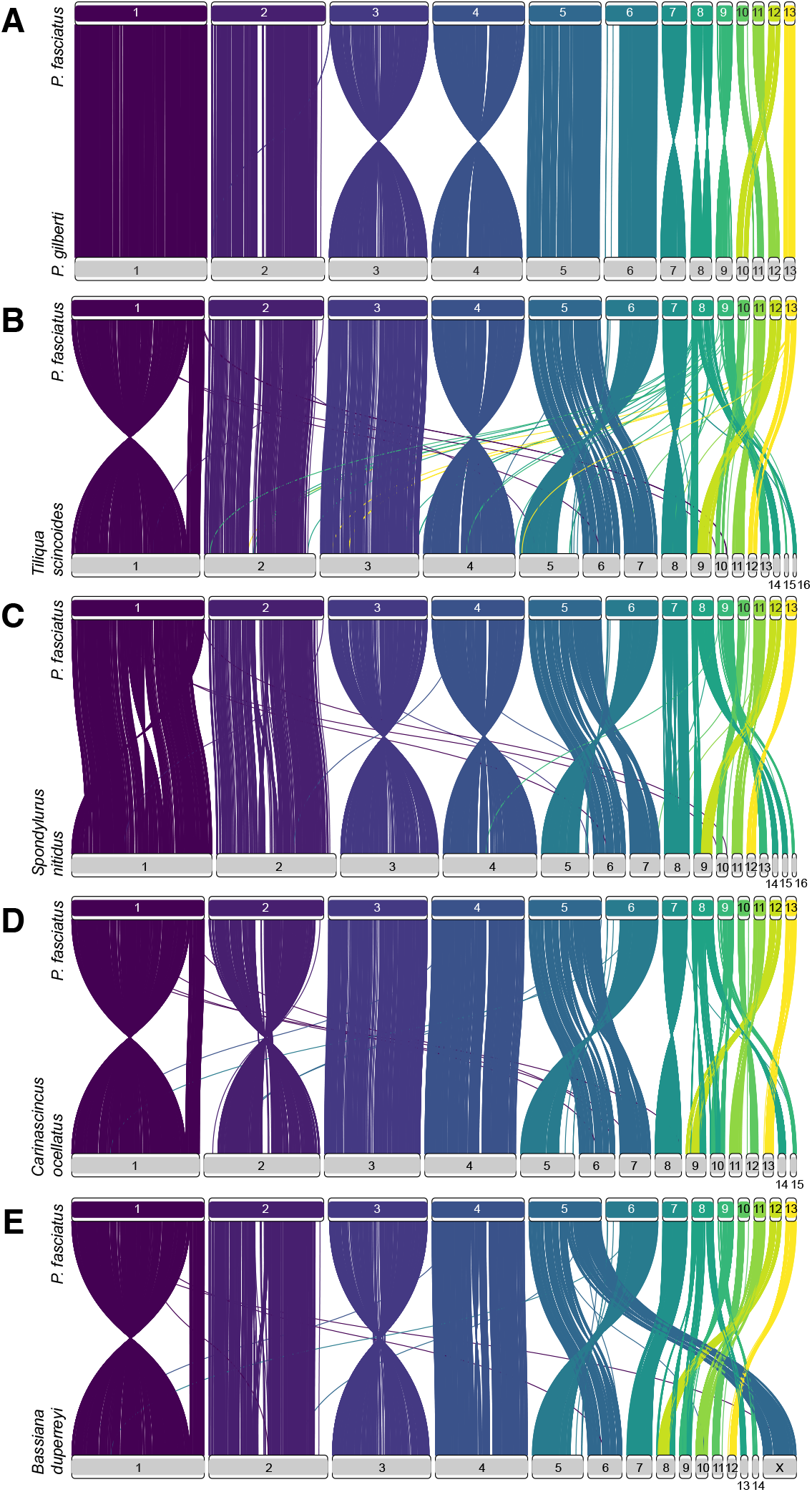
Synteny of BUSCO genes between *Plestiodon fasciatus* and other skinks. **A**. *P*. gilberti; **B**. *Tiliqua scincoides;* **C**. *Spondylurus nitidus*; **D**. *Carinascincus ocellatus;* **E**. *Bassiana duperreyi*.

### *P. fasciatus* and *P. gilberti*

Due to the cryptic, homogametic XY sex-determining system in many skinks, little is known about the position of sex-determining regions on the chromosomes; however, hypothetical sex chromosomes have been identified in *Bassiana duperryi* (Dissanayake et al., 2020; Hanrahan et al., 2025). Here, we show that a block of chromosome 5 is syntenic with the X chromosome in *B. duperreyi*, indicating a potential location for sex-determining regions in *P. fasciatus* (**Figure 3**). The same block of the fifth chromosome of *Plestiodon gilberti* is syntenic with the X chromosome of *B. duperreryi* (Richmond et al., 2025). In an attempt to further identify sex-linked regions, we used the *FindZX* pipeline (Sigeman et al., 2022) with two male and two female *P. fasciatus*; however, the results were inconclusive. Despite some preliminary work, more research is required to further classify sex-determination in skinks, ideally with population-level sampling of populations with numerous representatives from both sexes across Scincidae.

## Conclusion

We present a high-quality, telomere-to-telomere, chromosome-level annotated reference assembly of the North American common five-lined skink *Plestiodon fasciatus* (Linnaeus 1758), representing one of the most complete reference genomes (*rPleFas1*.*1*) to date of any species of Scincidae. We find that macrochromosome structure is conserved across the family, but there are common rearrangements of the microchromosomes, including a likely fusion in *P. fasciatus*, which has fewer microchromosomes than many other skink species. We also present insight into the content and evolution of transposable elements in skinks, which show remarkable conservation between species over the last ∼115Ma.

## Data availability

The data presented in the paper is available on NCBI (BioProject PRJNA1278702) and Figshare (https://figshare.com/projects/Telomere-to-telomere_reference_genome_of_the_common_five-lined_skink_Plestiodon_fasciatus_Squamata_Scincidae_/254723). The code for analyses can be found at https://github.com/jomhoff/Chromosome-level_genome_assembly and https://github.com/jomhoff/Genome-Annotation.

## Acknowledgments

The authors thank Megan Wallace of the American Museum of Natural History’s Institute of Comparative Genomics for her laboratory expertise, as well as Mac Mahacek, Steven Price, William Taylor, and Elijah Wessel for field assistance. The authors also thank the Michigan Department of Natural Resources and Kentucky Department of Fish and Wildlife. Finally, we thank two anonymous reviewers that improved the quality of this manuscript.

